# Are foundation species effects different than those of dominant species? A case study of ant assemblages in northeastern North American forests

**DOI:** 10.1101/062265

**Authors:** Sydne Record, Tempest McCabe, Benjamin Baiser, Aaron M. Ellison

## Abstract

Foundation species uniquely control associated biodiversity through non-trophic effects, whereas dominant species are locally abundant but are replaceable in ecological systems. Long-term data on ant assemblages at the Harvard Forest Hemlock Removal Experiment (HF-HeRE) and the Black Rock Future of Oak Forests Experiment (BRF-FOFE) provide insights into how ant assemblages change and reassemble following the loss of a foundation species (*Tsuga canadensis*) or a dominant genus (*Quercus*). At HF-HeRE, removal of *T. canadensis* trees resulted in taxonomic and functional shifts in ant assemblages relative to control stands. In contrast, ant assemblages at BRF-FOFE varied little regardless of whether oaks or non-oaks were removed from the canopy. Non-trophic effects of foundation species were stronger than indirect trophic effects on taxonomic and functional diversity of ant assemblages. In contrast, non-trophic effects of dominant species were weaker than indirect trophic effects on ant taxonomic diversity and some measures of ant functional diversity.

**Statement of authorship:** A.M. Ellison and S. Record conceived the study. A.M. Ellison, T. 17 McCabe, and S. Record collected field data. T. McCabe and S. Record did the taxonomic diversity analyses. B. Baiser and S. Record did the functional diversity analyses. All authors contributed to drafts of the manuscript.

**Data accessibility:** All data (*i.e.*, ant and trait) and R code are available from the Harvard Forest Data Archive (http://harvardforest.fas.harvard.edu/data-archive), datasets HF-118 (HF-HeRE) and HF-097 (BRF-FOFE). Nomenclature follows Bolton (2016); voucher specimens are stored at the Harvard Forest and at the Museum of Comparative Zoology.

## Introduction

Ecosystems with high biodiversity are hypothesized to be more resilient to changing environmental conditions than those with lower biodiversity because more species are available in the former to fill functional roles when species are lost (*i.e.*, the insurance hypothesis *sensu* Yachi & Loreau 1999). However, not all species are functionally equivalent, and the loss of particular ones, such as keystone predators (Paine 1966), dominant species (Whittaker 1965), or foundation species (Ellison *et al.* 2005) may have unexpectedly large or even irreversible system-wide impacts. In terrestrial environments, foundation species and dominant species tend to be trees: large, abundant, species that occupy basal positions in local food webs, and control ecosystem processes and dynamics principally through non-trophic interactions (Baiser *et al.* 2013). Foundation species are thought to be irreplaceable in terms of processes that they control, whereas dominant species are considered to be replaceable (Ellison *et al.* 2005).

The contrast between foundation species and dominant species can be seen clearly in the forests of eastern North America. *Tsuga canadensis* (L.) Carrière (eastern hemlock) is a foundation tree species in many of these forests (Ellison *et al.* 2005; 2014a); stands dominated by *T. canadensis* are different, both structurally and functionally, from stands dominated by other conifers or mixtures of various deciduous species. Hemlock-dominated stands are dark, cool, and moist; have acidic, nutrient-poor soils with slow rates of nutrient cycling (*e.g.,* Orwig *et al.* 2013), and are populated by generally species-poor assemblages of associated plants and animals (*e.g.*,Rohr *et al.* 2009; Sackett *et al.* 2011; Orwig *et al.* 2013).

On the other hand, many eastern North American forests are dominated in terms of numbers or biomass by one or more *Quercus* (oak) species (*e.g.*, Schuster *et al.* 2008). Unlike *T. canadensis,* however, oaks appear not to determine uniquely the structure and function of the forests they dominate. Although oak masts are important for certain organisms such as small mammals and ticks (Ostfeld *et al.* 1996; McShea *et al.* 2007), most core ecosystem processes of oak-dominated forests, including leaf-litter decomposition rates and associated soil nutrient dynamics (Polyakova & Billor 2007), root respiration (Levy-Varon *et al.* 2012), and net ecosystem exchange (Papale & Valentini 2003) are statistically indistinguishable from forests dominated by other deciduous species or from forests with no clear dominant species.

Many trees, including *T. canadensis* and *Quercus* spp. are threatened with functional loss by a myriad of native and nonnative insects and pathogens (Lovett *et al.* 2016). At the same time that we are mourning these impending losses (Foster 2014) and many researchers are working to control these insects and pathogens, we also have an unparalleled opportunity to study how forests respond to and reorganize after the loss of foundation or dominant species by assessing how their loss changes the taxonomic and functional biodiversity of associated species.

Here, we use two forest canopy-manipulation experiments to ask how ground-and soil-nesting ant assemblages change and reorganize after the experimental removal of either the foundation species *T. canadensis* or the dominant *Quercus* species. Ants are a particularly good taxon to use for these studies because they are abundant and widespread omnivores; are known to be responsive to local environmental conditions such as canopy cover, light availability, and soil temperature (Rescaso *et al.* 2014); modulate and control some soil ecosystem processes (Del Toro *et al.* 2012; Kendrick *et al.* 2015); and are well known both taxonomically and functionally in northeastern North America (Ellison *et al.* 2012; Del Toro *et al.* 2015).

We hypothesized that the loss of the foundation species would result in ant assemblages that differed in terms of species composition, relative abundance, and functional diversity from ant assemblages in control stands. In contrast we predicted that ant assemblages would be similarly structured (compositionally and functionally) in control stands and stands where oaks had been removed. Second, we hypothesized that when a foundation species was present that direct, non-trophic effects of the foundation species on ant assemblages would be stronger than indirect trophic effects (Fig. 1; Ellison *et al.* 2010). When a foundation species was absent and a dominant one was in its place, however, we expected trophic effects to have a greater influence on ant assemblages than the non-trophic effects (Fig. 1).

**Figure 1.**
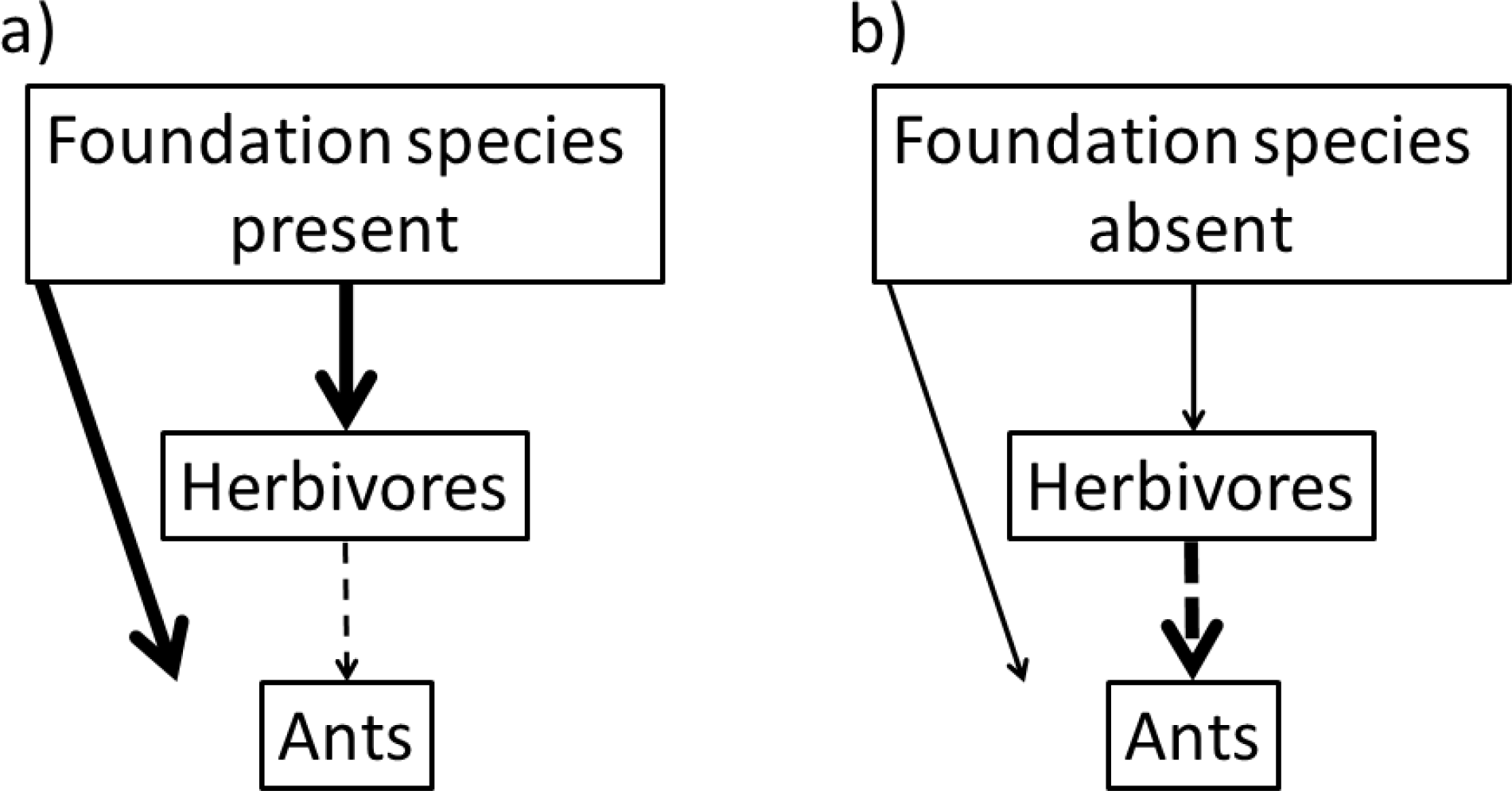
Conceptual framework illustrating hypothesized differences in the magnitude of non-trophic effects (solid arrows; *i.e.*, effects of foundation species) and indirect trophic effects (dashed arrows) on arthropod assemblages in the presence and absence of a foundation species. Arrow width indicates the strength of influence of the non-trophic and trophic effects.

## Materials and methods

### Harvard Forest – Hemlock Removal Experiment (HF-HeRE)

The Harvard Forest – Hemlock Removal Experiment (HF-HeRE) is a stand-level manipulation of *T. canadensis*-dominated forests located in the hemlock/hardwood/white pine transition zone of the temperature forest biome of northeastern North America. The experiment is located within the 121-ha Simes Tract at the Harvard Forest in Petersham, Massachusetts, USA (42.47° to 42.48° N, 72.21° W, 215-300m a.s.l.) HF-HeRE is a replicated and blocked Before-After-Control-Impact experiment. The experimental plots are 90 × 90 m in size; *T. canadensis* accounted for >70% of the basal area of the manipulated plots (see Ellison *et al.* 2010 for full details on the design and routine statistical analysis of HF-HeRE). Briefly, there are two replicates of each of four treatments, which are grouped into two blocks with equal representation of treatments per block. The ‘Valley’ block is sited in poorly-drained rolling terrain bordered on the north by a *Sphagnum*-dominated wetland, whereas the ‘Ridge’ block is along a forested ridge with well drained soils. Plots were identified in 2003 and pre-treatment data on plant and ant assemblages were collected for two years prior to any experimental manipulations. Soils at the Simes Tract are derived from glacial till and composed primarily of coarse-loamy, mixed, mesic Typic Dystrudepts in the Charlton Series (Giasson *et al.* 2013), and canopy trees within HF-HERE are 70-150 years old (Ellison *et al.* 2014).

There are two canopy-level manipulations in HF-HeRE. In the first, all *T. canadensis* individuals were girdled (as done by Yorks *et al.* 2003) to simulate physical impact of loss of hemlock caused by the nonnative hemlock woolly adelgid (*Adelges tsugae* Annand; Homoptera: Adelgidae). Girdling of all individuals and age classes of hemlocks (*i.e.*, seedlings through mature adults) occurred over 2 days in May 2005 using knives or chainsaws to cut through bark and cambium. Sap flow of girdled trees decreased immediately by 50%, and the trees died within 30 months (Ellison *et al.* 2010). This timeframe for hemlock death in the girdled plots was comparable to mortality times in response to adelgid infestation observed in the southeastern U.S., but somewhat faster than the 5-10 years observed routinely in the northeastern U.S. (McClure 1991). Dead trees were left standing in the girdled plots where they have slowly disintegrated and fallen, as in stands naturally infested by the adelgid. The volume of dead stumps and snags was comparable to the controls in pre-treatment years, but increased 2-3 orders of magnitude within two years post-girdling (Orwig *et al.* 2013).

In the second canopy manipulation, we cut and removed from the relevant plots all *T. canadensis* individuals > 20-cm diameter (measured at 1.3-m above ground) along with other select merchantable trees (*e.g., Pinus strobus* L., *Quercus rubra* L.) to simulate the pre-emptive salvage logging done by people in anticipation of the arrival of the adelgid (Foster and Orwig 2006). Logging was done between February and April 2005 when the soil was frozen. Logging removed 60-70% of the basal area in these plots; slash (*i.e.*, small branches and damaged or rotting boles) were left on site as is typical in local timber-harvesting operations (Ellison *et al.* 2010).

There also are two controls in each block at HF-HeRE: intact hemlock (~70% basal area hemlock) and intact mixed hardwood stands. When the experiment was sited in 2003, the adelgid was absent from the region. By 2009, however, the adelgid was observed at low densities in the hemlock control plots, and it was widespread in these plots by 2010. The hardwood controls represent the anticipated future of stands in this region following hemlock decline and contain mixed young hardwoods (predominantly *Betula lenta* L. and *Acer rubrum* L. <50 years old) and small *T. canadensis* and *P. strobus* individuals (Ellison *et al.* 2010). To ensure that any discernible differences in ant assemblages were due to the experimental manipulations (as opposed to known environmental heterogeneity), within each block, the hemlock removal treatments and control plots were sited within 100 m of one another and occupy similar topography, aspect, and soil type. Finally, in 2012, 15 × 30 × 2.5-m exclosure fences were erected in the center of each canopy manipulation or control plot to examine additional effects of large browsers (moose: *Alces alces* L. and White-tailed deer *Odocoileus virginianus* Zimmerman) on successional processes (Faison *et al.* 2016).

To document temporal changes in ant assemblages within the eight HF-HeRE plots, ants were sampled annually from 2003-2015 using pitfall traps, baits, litter samples, and hand collections (the ALL protocol: Agosti & Alonso 2000) within a permanent 10 × 10-m grid with 25 evenly-spaced sampling points located near the center of each canopy-manipulation plot (sampling details in Sackett *et al.* 2011). In 2012, following the installation of the large herbivore exclosures, we added an entire additional sampling grid for ants within each exclosure, thus doubling the total sample size.

### Black Rock Forest – Future of Oak Forests Experiment (BRF-FOFE)

The Black Rock Forest – Future of Oak Forests Experiment (BRF-FOFE) is a canopy-level manipulation of oak and non-oak trees. Black Rock Forest is located in the Hudson Highlands near Cornwall, New York (Ellison *et al.* 2007) and BRF-FOFE is located on the north slope of Black Rock Mountain (400 m a.s.l; 41.45°N, 74.01°W) within a “hardwood slope” (Tryon 1930) or “red oak association” (Raup 1938). Like HF-HeRE, the BRF-FOFE is a replicated and blocked Before-After-Control-Impact experiment. The experimental plots are 75 × 75 m. There are three replicates of each of four treatments, which are grouped into blocks with equal representation of treatments per block. The blocks represent slope position (lower, middle, upper). Upper slope plots are steeper and have drier soils than the lower slope plots (15-16% and 24% soil water content, respectively; Falxa-Raymond *et al.* 2012). Plots were sited in 2006 and pre-treatment data on ant assemblages were collected for two years prior to experimental manipulations. The glacially derived soils at Black Rock Forest are of the Chatfield and Rockaway series (Denny 1938, Ross 1958). *Quercus rubra* and *Q. prinus* L. make up ~67% of the canopy; *Acer rubrum* dominates the understory (Schuster *et al.* 2008). Other non-oak canopy trees in decreasing order of prevalence include: *A. rubrum* (28%), *Nyssa sylvatica* Marsh. (22%), *Betula lenta* (20%), and *A. saccharum* L. (16%) (Falxa-Raymond *et al.* 2012). *Tsuga canadensis* is absent from the experimental site.

There are four experimental treatments in BRF-FOFE: all oaks girdled (OG; girdling 74-78% of total plot basal area (BA)); half of the oaks girdled (O50; 15-37% of BA); all non-oaks girdled (NO; 15-37% of BA); and control (no trees girdled; 0% of BA). The intent of the girdling treatment was to simulate effects of pathogens (*e.g.*, Sudden Oak Death: *Phytopthera ramorum*

Werres, de Cock & Man in’t Veld) or defoliating insects such as gypsy moth (*Lymantria dispar* (L.)). Although Sudden Oak Death is not yet epidemic in the eastern U.S. (as it is in northern California and southern Oregon), eastern U.S. nurseries have documented infected horticultural stock arriving from western sources and there is concern that it could become established and virulent in eastern North American forests (Grünwald *et al.* 2012).

Between June 27 and July 9, 2008, chainsaws were used in each plot to girdle trees > 2.54-cm diameter by making incisions up to 5-cm deep so as to cut through the bark, phloem, and cambium around the entire circumference of each tree at 1.3-m above ground. The many deer at BRF (≈7/km^2^ at the start of the experiment) had browsed most of the understory plants and < 3% of the trees were < 2.54 cm in diameter and so were not girdled. Unstable girdled trees, typically those 2.5 – 7.5-cm in diameter, were felled for researcher safety. One year after girdling, in the summer of 2009, roughly twice as many oak trees leafed out and re-sprouted in the 050 plots (8% and 27%, respectively) as in the OG plots (15% and 46%, respectively). After girdling, over twice as many non-oaks leafed out and re-sprouted (23% and 69%, respectively) than oaks (10% and 33%, respectively; Levy-Varon *et al.* 2012). Trees that survived girdling were re-treated as needed in subsequent years. In addition to these treatments, each plot contained a 10 × 20 × 3-m exclosure fence to keep out deer and examine effects of browsing on successional trajectories.

To document temporal changes in ant assemblages within the twelve BRF-FOFE plots, ants were sampled in 2006 using the ALL protocol (pitfalls, baits, litter sifting, hand collection; Agosti & Alonso 2000) and then with hand collections from nests for 1 person-hour and litter sifting biennially from 2007-2011 and again in 2015 (Ellison et al. 2007). Hand sampling within the deer exclosures was done for 10 person-minutes (time spent sampling within the 200-m^2^exclosures was downscaled from the time spent sampling the rest of the 5625-m plots). Specimens were placed directly into 95% EtOH in the field, and subsequently identified at Harvard Forest.

### Assessing taxonomic composition in ant assemblages over time

Estimates from pitfall traps, litter samples, or baits may overestimate abundance of ants if they happen to occur nearby colonies with actively foraging workers, so we conservatively estimated abundance based on incidences of species from baits, litter samples, or pitfall traps (Gotelli *et al.* 2011). Each hand-collected sample was from a separate nest, and so each was treated as an incidence when estimating abundance. Species accumulation curves and rarefaction analyses for estimates of Hill numbers for species richness (^0^*q*), Shannon diversity (^1^*q*), and Simpson’s diversity (^2^*q*) of ant data (Jost 2007) assessed sampling efficacy for each experiment using the iNEXT package v.2.0.5 of R statistical software v.3.2.3 (Hsieh *et al.* 2016; R Core Team 2015).

We used non-parametric multivariate analyses of variance (npMANOVA) and covariance (npMANCOVA) to assess variation in ant assemblages prior to manipulation (*i.e.*, pre-treatment) and over time among treatments post-treatment for both experiments. Prior to running multivariate analyses, data were screened for multivariate outliers and to check for nonsignificant multivariate dispersion of factors using the ‘betadisper’ function of the R vegan package (Oksanen *et al.* 2016). The response variable was the Bray-Curtis distance matrix of pairwise distances between species’ relative frequencies. In all analyses, block entered the model as a random factor; all other factors were considered fixed effects. In the post-canopy treatment npMANOVA for HF-HeRE, the data were split into pre-and post-adelgid infestation temporal strata (Appendix S1, Table S1). Since large herbivore exclosures in the HF-HeRE were installed several years after the canopy manipulations and after the adelgid infested the plots, the herbivore exclosure analysis for HF-HeRE does not include temporal stratum (pre/post-adelgid) as a factor. In the BRF-FOFE analysis, year entered the model as a covariate rather than as a factor and we used a canopy treatment x year interaction to explore lagged effects of the canopy manipulations. Estimates of F-statistics were calculated based on 5,000 permutations using the ‘adonis’ function from the vegan package v. 2.4 of R v.3.2.3. For both experiments, pairwise comparisons were used to compare differences between treatments.

To illustrate compositional differences in ant assemblages between canopy and exclosure treatments, ordination plots were generated based on non-metric multidimensional scaling (NMDS) using 200 random starts in search of a stable solution with the ‘metaMDS’, ‘ordiellipse’, and ‘ordiplot’ functions of the R vegan package. When fitting the NMDS ordinations, scree plots were examined to determine the optimal number of dimensions based on the minimum number of dimensions that resulted in <0.05% stress.

### Assessing changes in functional composition over time

To assess changes in functional composition over time in ant assemblages, we analyzed functional beta-diversity based on a species-by-trait matrix that included twelve continuous, binary, and categorical traits. These traits reflect major ant-mediated ecosystem processes: soil movement, decomposition, seed dispersal, and regulation of invertebrate and plant community structure (Del Toro *et al.* 2015). We used trait definitions from Del Toro *et al.* (2015) and filled in missing species’ data with information from Ellison *et al.* (2012), www.antweb.org, and www.antwiki.org.

We calculated a species-by-species distance matrix from the species-by-trait matrix using the Gower coefficient (Gower 1971), which can accommodate ordinal, nominal, and binary data. Two undescribed species (*Leptothorax sp.* and *Myrmica* sp. with species codes AF-can and AF-smi, respectively, in Ellison *et al.* 2012) were not included in the analysis as data on their traits were unavailable. Using this distance matrix, we calculated two commonly used measures of functional beta-diversity (Ricotta & Burrascano 2008; Swenson 2014): the abundance-weighted mean pairwise distance (*D_pw_*) and abundance-weighted nearest neighbor distance (D_nn_).We calculated *D_pw_* and *D_nn_* using the ‘comdist’ and ‘comdistnt’ functions, respectively, from the picante packagev.1.6-2 in R v.2.3.3 (Kembel *et al.* 2015).

Many measures of functional beta-diversity (including *D_pw_*) are highly correlated with measures of taxonomic beta-diversity (*e.g.*, pairwise Bray-Curtis similarity on species composition) (Swenson 2014). To determine if functional turnover was greater (or less) than expected based on taxonomic turnover alone, we used null models to calculate standardized effect sizes (S.E.S.) for each metric (Swenson 2014). We constructed our null model by shuffling the species names in the species-by-trait matrix, calculating a distance matrix from the shuffled trait matrix, and then calculating the beta-diversity metric. We did this randomization 999 times for each of the two metrics, then calculated the S.E.S. as 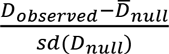 for the two diversity indices *D_pw_* and *D_nn_*. Large S.E.S. values (> |2|) indicated more (> 2) or less (< −2) functional beta-diversity than expected by chance alone. Because we were specifically interested in functional change within each treatment plot through time relative to the pre-treatment conditions, we examined further those pairwise *D_pw_, D_nn_*, and S.E.S values.

Differences among canopy and deer exclosure treatments in *D_pw_, D_nn_*, and S.E.S values were tested using generalized linear mixed models (GLMMs) in which the predictor variables and number of tests for each experiment matched the post-treatment analyses outlined in Appendix S1, Table S1. Block entered as a random effect, whereas other factors entered the GLMMs as fixed effects. Residual plots of each GLMM ensured model assumptions were met. Given the number of significance tests performed in this study, a Holm correction (Holm 1979) to control for Type I error was performed on all test statistics generated in the analyses with the ‘p.adjust’ function of the ‘stats’ package in R version 3.2.3.

### Data and specimen availability

All data (*i.e.*, ant and trait) and R code are available from the Harvard Forest Data Archive (http://harvardforest.fas.harvard.edu/data-archive), datasets HF-118 (HF-HeRE) and HF-097 (BRF-FOFE). Nomenclature follows Bolton (2016); voucher specimens are stored at the Harvard Forest and at the Museum of Comparative Zoology.

## Results

At HF-HeRE, we accumulated 47 ant species (2941 incidence records) from 2003-2015, but species accumulation has not yet reached an asymptote (Appendix S2). In contrast, we accumulated 48 ant species (1882 incidence records) at BRF-FOFE from 2006-2015 and the species accumulation curve reached an asymptote in 2015 (Appendix S2). Rarefaction curves for both sites indicated sufficient sampling coverage for estimation of Shannon and Simpson diversity indices, but indicated that more samples would help to better determine richness, especially at HF-HeRE (Fig. 2). Rarefaction curves also revealed that estimates of effective species richness, Shannon, and Simpson’s diversity at HF-HeRE consistently were highest for the logged treatment, intermediate for the girdled and hardwood treatments, and lowest for the girdled treatment (Fig. 2). At BRF-FOFE, estimates of effective species richness and Shannon diversity of ants was highest in the 100% oak-girdled treatment, intermediate in the control and non-oak girdled treatment, and lowest in the 50% oak-girdled treatments (Fig. 2). Differences among treatments in effective Simpson’s diversity were negligible across treatments at BRF-FOFE.

**Figure 2.**
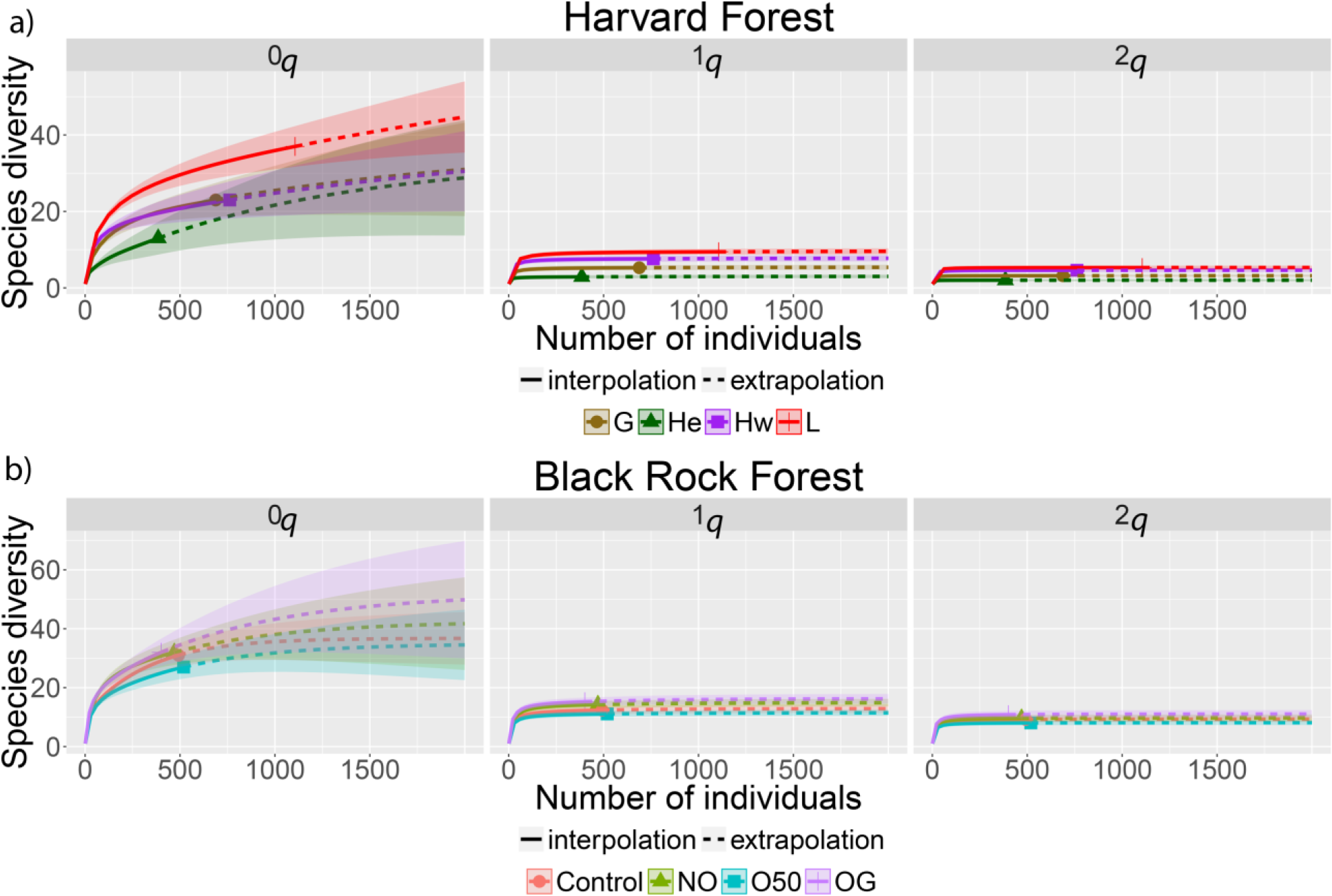
Sample size based rarefaction/extrapolation curves of ant incidence frequency data from collections made in the Harvard Forest Hemlock Removal Experiment (HF-HeRE) from 2003-2015 and in the Black Rock Forest Future of Oak Forests Experiment (BRF-FOFE) from 2006-2015. Colors indicate treatments: Solid lines are observed estimated Hill numbers for species richness (^0^*q*), Shannon diversity (^1^*q*), and Simpson’s diversity (^2^*q*), whereas dashed lines are extrapolations of the curves.

### Taxonomic composition in ant assemblages over time

#### Harvard Forest Hemlock Removal Experiment

Prior to canopy manipulations at HF-HeRE, there were no significant differences in the composition of ant assemblages across plots selected for canopy treatments (NPMANOVA: *F*_2_,_9_ = 0.7, *P* = 1.0). Following canopy manipulations there were significant differences in composition among canopy treatments (NPMANOVA: *F*_3_,_111_ = 11, *P* = 0.02; axis NMDS2 of Fig. 3a) and temporal strata (NPMANOVA: *F*_1_,_111_ = 175, *P* = 0.02; axis NMDS1 of Fig. 3a), but there was no effect of canopy treatment nested within temporal stratum (NPMANOVA: *F*_3_, _111_ = 0.76, *P* =1.0). All pairwise comparisons between canopy treatments indicated significant differences (*P* < 0.05) in the composition of ant assemblages, except for the pairwise comparison between the girdled and hardwood control treatments (Fig. 3a). For functional beta-diversity, there were significant canopy treatment effects for all measures (*i.e.*, the abundance-weighted mean pairwise distance, *D_pw_*, nearest neighbor distance, *D_nn_* and their standardized counterparts SES *D_pw_* and SES *D_nn_*; *P* < 0.05). The ant assemblage in the hemlock control treatment was functionally more similar to its pre-treatment assemblage than the other three treatments were to their pre-treatment assemblages according to SES *D_pw_* (Fig. 4a). The same was true for the hemlock treatments when compared to the hardwood and logged treatments with the *D_pw_* (Fig. 4b). According to the SES *D_nn_* and *D_nn_* measures, the ant assemblage in the hemlock control treatment was functionally no more different from its pre-treatment assemblage than the hardwood and logged treatments were to their pre-treatment assemblages (Fig. 4C, D). However, SES *D_nn_* and *D_nn_* showed that the ant assemblage in the hemlock control treatment was functionally less similar to its pre-treatment assemblage than the girdled treatment was to its pretreatment assemblage (Fig. 4C, D). Finally, the composition (Fig. 5b) and functional diversity (Fig. 6a-d) of ant assemblages were similar regardless of whether they were collected from within or outside of herbivore exclosures at HF-HeRE.

**Figure 3.**
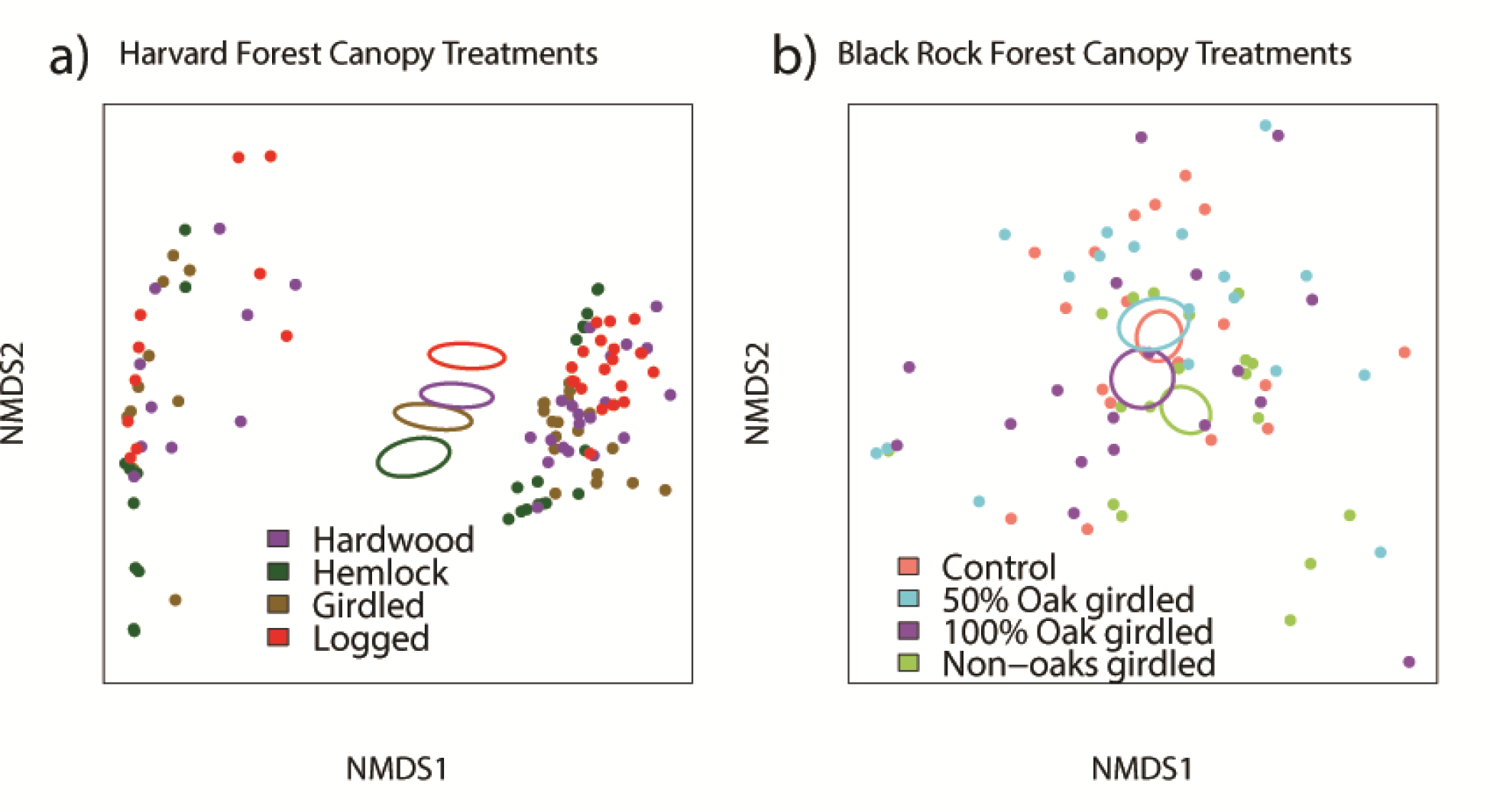
Non-metric multidimensional scaling (NMDS) ordination plot illustrating differences among sampling plots plotted in species space by canopy treatment at a) HF-HeRE and b) BRF-FOFE. Although only the first two axes (k=2) of the NMDS ordinations are shown here, HF-HeRE NMDS was generated with *k*=7 (linear fit *R*^2^ = 0.974; non-metric fit *R^2^* = 0.998) and the BRF-FOFE NMDS with k=8 (linear fit *R^2^* = 0.981; non-metric fit *R^2^* = 0.998). Ovals represent standard errors of the weighted average of NMDS scores for each treatment. For the HF-HeRE NMDS (a), NMDS1 separates the samples temporally with samples from before adelgid infestation in 2009 to the left and post-adelgid infestation to the right, whereas NMDS2 separates the samples by canopy treatment.

**Figure 4.**
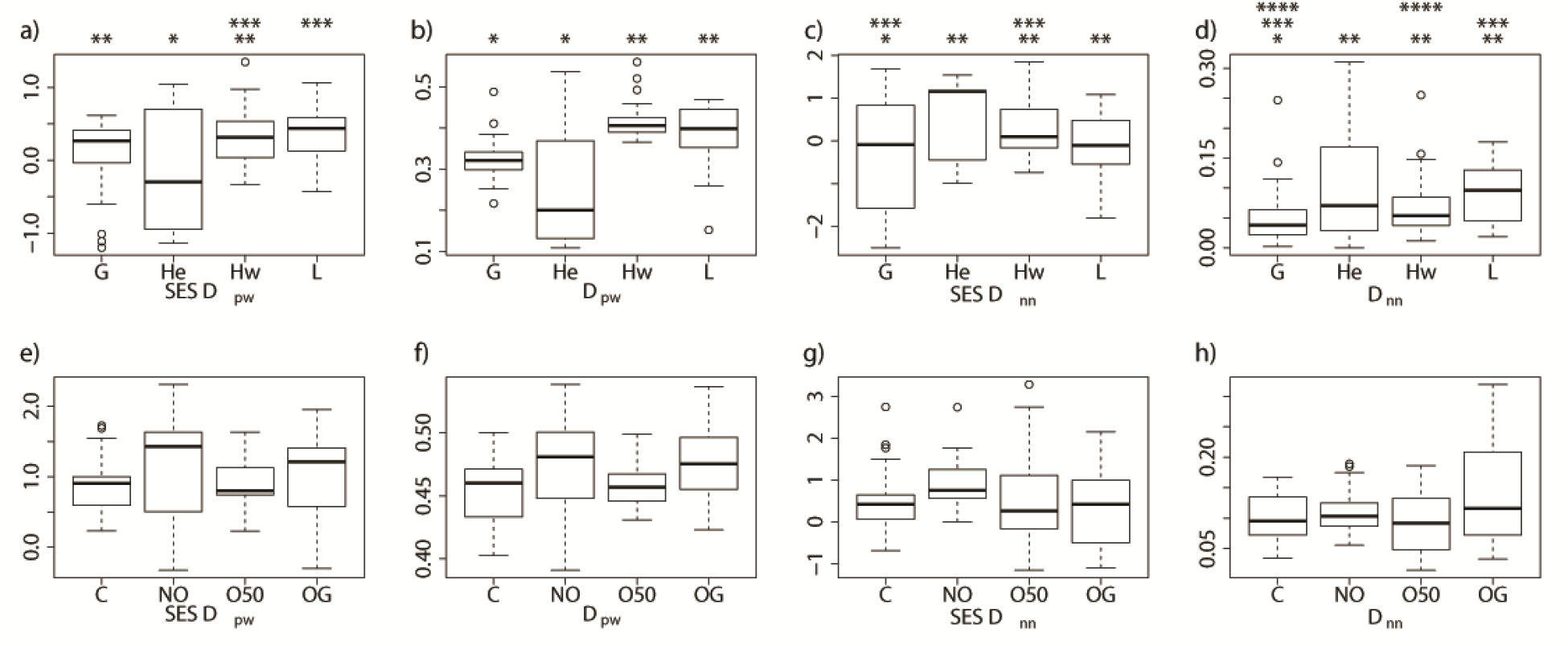
Boxplots illustrating differences between canopy treatments at HF-HeRE (a-e) and BRF-FOFE (f-j) in terms of functional diversity (*i.e.*, the abundance-weighted mean pairwise distance based on standardized effect size (SES *D_pw_;* a, e) and raw (*D_pw_*; b, f) estimates and the abundance-weighted nearest neighbor distance based on standardized effect size (SES *D_nn_*; c, g) and raw (*D_nn_*; d, h) estimates). Asterisks above the boxplots indicate significant differences (*P* < 0.05) for post-hoc pairwise comparisons between canopy treatments. If no asterisks are shown above a boxplot, then there were no significant effects of canopy treatment. Canopy treatments are labeled on the x-axis as follows for HF-HeRE (a-d): girdled (G), hemlock control (He), hardwood (Hw), and logged (L). Canopy treatments are labeled on the x-axis as follows for BRF-FOFE (e-h): control, no trees girdled (C); non-oaks girdled (NO); 50% oaks girdled (O50); and 100% oaks girdled (OG).

#### Black Rock Forest Future of Oak Experiment

Prior to canopy manipulations at BRF-FOFE, there were no significant differences in the composition of ant assemblages across plots that were allotted to the canopy manipulations (F_3_,_20_ = 1.1, *P* = 1.0). Neither canopy manipulation nor the canopy by year interaction influenced taxonomic or functional diversity at BRF-FOFE (Figs. 4, 5e-h), and the functional diversity of ant assemblages in the pre-and post-treatment plots within each treatment were not different across treatments regardless of the measure used (Fig. 4e-h). Furthermore, there were no significant effects due to canopy, year or their interaction (*P* > 0.05, Fig. 4e-h).

Ant assemblage composition at BRF-FOFE was influenced by excluding herbivores (Fig. 5b), but not by the exclosure × canopy manipulation interaction. Ant assemblages that were on the exterior of the herbivore exclosure were functionally more similar to their pre-exclosure assemblages than those within the exclosure were to their pre-exclosure assemblages (*D_nn_:* F_1_,_58_ = 50, *P* < 0.0001; Fig. 6h). However, all other functional beta-diversity measures indicated no significant effects of the exclosure treatment (P > 0.05; Fig. 6e-h). Similarly, none of the functional beta-diversity metrics indicated an effect of an exclosure × canopy treatment interaction at BRF-FOFE (*P* > 0.05).

**Figure 5.**
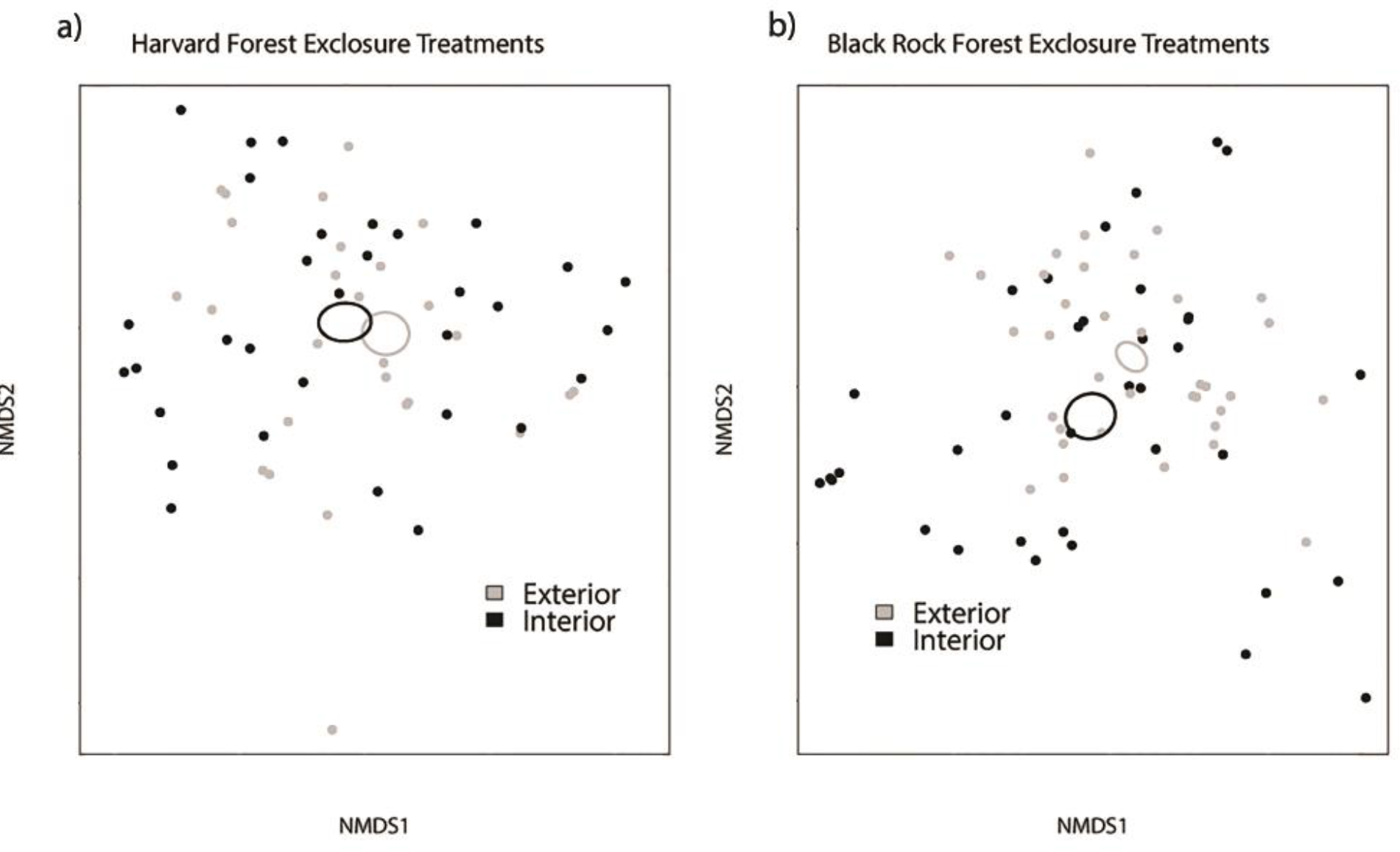
Non-metric multidimensional scaling (NMDS) ordination plot illustrating differences among sampling plots plotted in species space by herbivore exclosure treatment in the a) HF-HeRE and b) BRF-FOFE. Although only the first two axes (k=2) of the NMDS ordinations are shown here, HF-HeRE NMDS was generated with *k=8* (linear fit *R^2^* = 0.984; non-metric fit *R^2^* = 0.998) and the BRF-FOFE NMDS with k=8 (linear fit *R^2^* = 0.974; non-metric fit *R^2^* = 0.998). The composition of (a) ant assemblages was similar regardless of whether they were collected from within or outside of herbivore exclosures at HF-HeRE (NPMANOVA: F_155_ = 1.3, *P* = 1.0) and there was no exclosure × canopy treatment interaction (NPMANOVA: F_3_,_55_ = 1.8, *P* = 1.0). At BRF-FOFE (b), there were main effects of the herbivore exclosure treatment on the composition of ant assemblages (NPMANCOVA: F_1_,_60_ = 5.1, *P* = 0.02; Figs. 6b), but there was no herbivore × canopy manipulation interaction (NPMANCOVA: F_3_,_60_ = 0.88, *P* = 1.0). Ovals represent standard errors of the weighted average of NMDS scores for each treatment.

**Figure 6.**
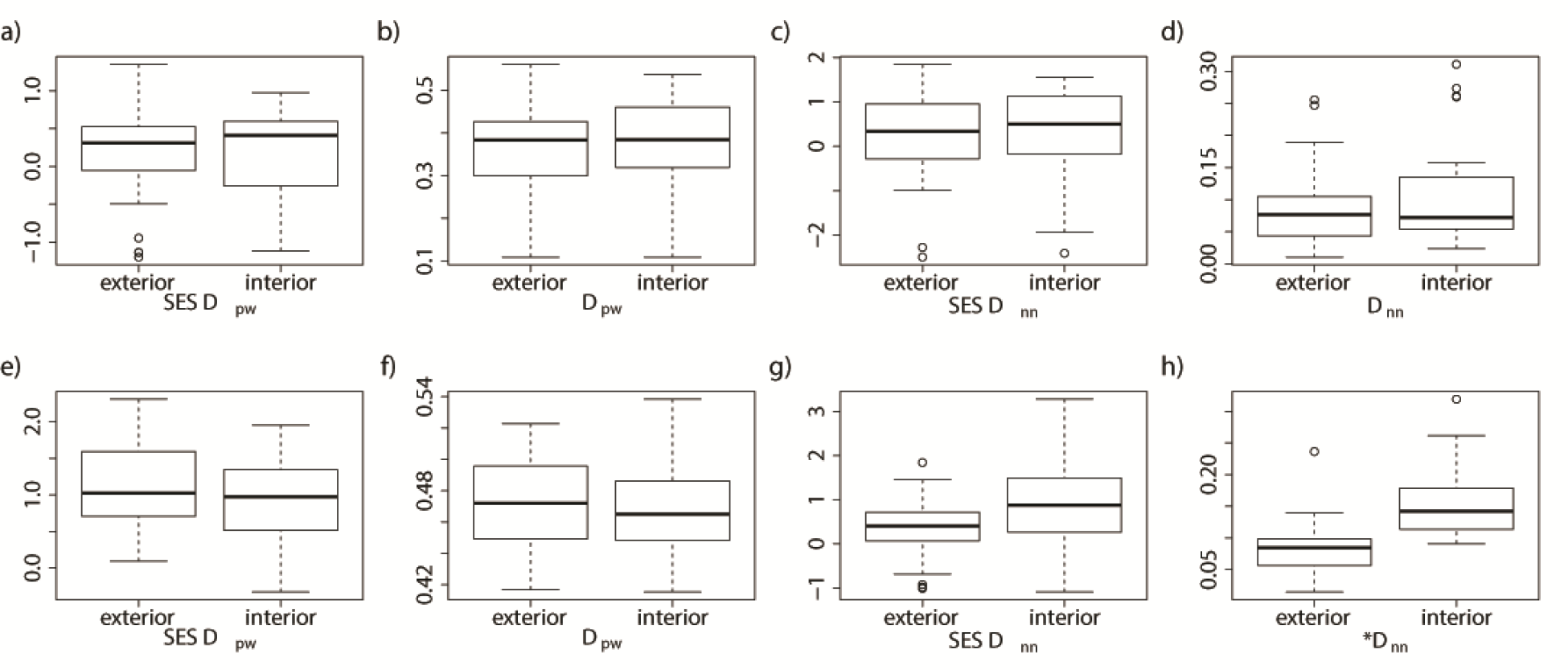
Boxplots illustrating differences between exclosure treatments in the HF-HeRE (a-e) and BRF-FOFE (f-j) in terms of functional diversity (*i.e.*, the abundance-weighted mean pairwise distance based on standardized effect size (SES *D_pw_*; a, e) and raw (*D_pw_*; b, f) estimates and the abundance-weighted nearest neighbor distance based on standardized effect size (SES *D_nn_*; c, g) and raw (*D_nn_*; d, h) estimates). Asterisks above a boxplot indicate significant effects of exclosure treatments. Canopy treatments are labeled on the x-axis as follows for HF-HeRE (a-d): girdled (G), hemlock control (He), hardwood (Hw), and logged (L). Canopy treatments are labeled on the x-axis as follows for BRF-FOFE (e-h): control, no trees girdled (C); non-oaks girdled (NO); 50% oaks girdled (O50); and 100% oaks girdled (OG).

## Discussion

The insurance hypothesis (Yachi & Loreau 1999) predicts that higher biodiversity will maintain ecosystem function in the face of unstable environments because more species are available to fill functional roles when species are lost. Although there is mounting evidence for the insurance hypothesis (*e.g.*, Gross *et al.* 2014), the role of foundation species should not be overlooked in this context with regards to maintaining biodiversity and ecosystem function. Given that foundation species are considered to be unique in terms of the processes that they control (Ellison *et al.* 2005) it is important to distinguish them from replaceable, dominant species and also from other factors determining structure and function of ecosystems (*e.g.*, trophic dynamics). The results reported here are the first that can be used to reveal differing effects of dominant and foundational tree species. Simultaneously, the large herbivore exclosure treatments in both experiments enable us to disentangle non-trophic effects of foundation species from trophic effects of large herbivores (see also Baiser *et al.* 2013), another important driver of biodiversity changes in temperate forests.

We hypothesized that the loss of a foundation species would result in ant assemblages that differed in species composition, relative abundance, and functional diversity from ant assemblages in control stands. Data from HF-HeRE provide strong support for this first hypothesis in terms of taxonomic composition. Our analysis of functional diversity showed that: ant assemblages in intact hemlock plots were functionally more similar to their pre-treatment assemblages than were those in the hardwood control and logged plots for SES *D_pw_* and *D_pw_* and girdled plots for SES *D_pw_* (Fig. 3a, b). This result supported our hypothesis that the adelgid has already caused some environmental changes (*e.g.*, increased light levels; Kendrick *et al.* 2015) in the hemlock stands resulting in functional change of ant assemblages. However, the nearest neighbor set of metrics did not corroborate these findings. Differences between pairwise measures and nearest neighbor measures are not uncommon (*e.g.*, Liu *et al.* 2016). Based on this discrepancy, overall we interpret our functional beta-diversity results as modest support for our hypothesis.

We also observed that the hemlock control plots had lower species richness than all other treatments. It is important to note that rarefaction analyses (Fig. 2) indicated that more intensive sampling in this study could have resulted in diminished differences between all treatments except for the logged treatment for estimates of species richness, but diversity estimates accounting for relative frequencies of species occurrence (*i.e.*, Shannon and Simpson’s diversity indices) likely would not have changed with increased sampling effort. Similar to previous research on ants within the HF-HeRE (Sackett *et al.* 2011; Kendrick *et al.* 2015), we found that hemlock control plots were largely dominated by *Aphaenogaster picea* and *A. fulva,* but lacked the *Formica* spp. and many of the *Lasius* spp. that colonized all the manipulated plots (see Harvard Forest data archives for full species lists by treatment).

Functionally, the hemlock control plots harbored fewer socially parasitic or behaviorally submissive species than the other canopy manipulation treatments. The logged and girdled plots with their greater amounts of coarse woody debris (Orwig *et al.* 2013) had the highest frequencies of occurrences of larger-bodied carpenter ants (*Camponotus* spp.). Rohr *et al.* (2009) compared arthropod communities (Arachnida, Chilopoda, Diplopoda, Insecta), including ants, in intact hemlock and hardwood stands within Shenandoah National Park in the mid-Atlantic
Appalachian Mountains of the eastern U.S. and also found that arthropod abundance and richness was lower in hemlock compared to hardwood stands, but that functionally the two forest types were similar.

At BRF-FOFE, our data supported the alternative hypothesis that ant assemblages would be compositionally and functionally similar whether or not oaks were removed (Figs. 4b, 5d-h). Together with the data from HF-HeRE, these results provide additional support for the hypothesis that *T. canadensis* is a foundation species for ant assemblages, whereas oaks are not. However, other species and ecosystem functions may respond differently to oak loss. For example, Bray (2015) suggested that oaks function as a foundation species for mites, collembola, and some arachnids at BRF-FOFE. Levy-Varon (2014) found that soil respiration in 100% oak girdled plots declined much more rapidly than in the control, non-oak girdled, or 50% oak girdled plots; these differences persisted from two weeks to two years post-treatment. Potential loss of oaks and concomitant increases in other hardwood species could have broader effects on forest dynamics that will depend on newly-established species, their interactions with pests and pathogens (Spaulding & Rieske 2011), and functional characteristics (Falxa-Raymond *et al.* 2012).

Our data also allowed us to test the hypothesis that direct, non-trophic effects of foundation species on ant assemblages would be stronger than indirect trophic effects of browsing mammals (Fig. 1; see also Ellison *et al.* 2010). At HF-HeRE we found evidence both for and against this hypothesis. The observations that herbivore exclosures did not influence ant taxonomic or functional diversity supported this model, but the lack of significant effects of canopy manipulation × exclosure on functional or taxonomic diversity of ants did not support it.

When a foundation species was absent and a dominant one was in its place, we expected trophic effects to have a greater influence on ant assemblages than the non-trophic effects. This hypothesis was supported strongly for taxonomic diversity but not for functional diversity at BRF-FOFE. The large browsers in New York and Massachusetts are moose and white-tailed deer. At the start of BRF-FOFE, deer densities were higher (7.3 km^−2^) than at Harvard Forest (4.2-5.7 km^−2^), but over the course of the study increased culling of the deer population at BRF has brought population densities down to comparable levels with New England (Personal communication W.S.F. Schuster). The relatively low densities of herbivores in New England and the fluctuating populations of deer in Black Rock Forest over the course of the study complicate our ability to disentangle non-trophic effects of foundation species and indirect trophic effects of browsers on local ant assemblages.

Data from two long-term hectare-scale experiments in eastern North America at Harvard and Black Rock Forests illustrate how ant assemblages reassemble after the loss of a foundation tree species versus a dominant canopy species, respectively. The loss of a foundation species at the HF-HeRE resulted in taxonomic and functional changes of ant assemblages in response to direct effects of the foundation species versus indirect trophic effects. The loss of a dominant species at the BRF-FOFE did not result in significant taxonomic or functional changes in ant assemblages, and trophic effects outweighed effects of canopy tree species loss whether oaks or non-oaks. These findings illustrate the importance of distinguishing between the roles of irreplaceable foundation versus replaceable dominant species.

## Acknowledgements

The group’s work on hemlock forests is part of the Harvard Forest Long Term Ecological Research (LTER) Program supported by grants 06-20443 and 12-37491 from the U. S. National Science Foundation. T.D.M. was supported by the Harvard Forest Research Experiences for Undergraduates Program and Bryn Mawr College’s Leadership Innovation and Liberal Arts Center. This paper is a contribution of the Harvard Forest LTER Program.

## Supporting Material

**Appendix S1**. Additional methodological details.

**Appendix S2**. Species accumulations curves for each treatment for HF-HeRE and BRF-FOFE.

